# Global patterns in the evolutionary relations between seed mass and germination traits

**DOI:** 10.1101/2024.04.16.589747

**Authors:** Keyvan Maleki, Filip Vandelook, Arne Saatkamp, Kourosh Maleki, Siavash Heshmati, Elias Soltani

## Abstract

During stressful climatic periods, plant populations face significant challenges, especially during germination and seedling establishment. Theoretical studies present conflicting findings regarding the relationship between seed size and germination rate. Some analyses suggest that species with larger seeds should exhibit rapid germination, while others point to faster germination in smaller seeds. This discrepancy can, for example, be attributed to the higher vulnerability of larger seeds to post-dispersal seed predation. To assess the correlation between seed mass and seed germination characteristics at a global scale, we performed a rigorous meta-analysis of published data sources, covering a wide range of germination traits across 1877 species with diverse dormancy types and global distribution. Additionally, we investigated the potential relationship between seed mass and dormancy level. Our findings revealed contrasting responses of germination traits to seed mass, suggesting complex eco-evolutionary correlations among these traits. Trade-offs and corelated evolution likely played an important role in the evolutionary history of seed mass, with seed mass displaying notable trait conservation across the phylogeny. Interestingly, our study demonstrated that seed dormancy exhibited trade-offs with post-germination traits and seed mass. The meta-analysis demonstrated that any changes in the relationship between seed mass and dormancy is dependent on dormancy types with physiological dormancy showing lower seed mass while other dormancy types were has a higher seed mass. These results underscore the critical role of seed mass in shaping plant performance and provide valuable insights into plant trait evolution. Our findings clearly indicate that the hypothesis of larger seeds conferring advantages in both pre- and post-germination traits might not be empirically true in all cases as theoretical studies suggest and that this relationship is complex and varies among species and growth forms. We interpret these results within the context of germination strategies and correlated evolution between seed dormancy and seed mass. Considering these traits in future analyses of plant germination strategies and species distributions is crucial. Factors such as lifespan, seed mass, and germination-related traits should be carefully considered to gain a comprehensive understanding of plant adaptation to challenging environmental conditions.

## Introduction

Climate-related uncertainty has been a motivating factor for ecologists to focus on the understanding of functional traits and their interaction with life history transitions, which are important determinants of plant persistence in varying environments (Buckley & Kingsolver, 2012; De Frenne et al., 2013; Gremer et al., 2020a; Maleki et al., 2024). Seed dormancy is a strategy ensuring plant survival in an unpredictable environment, by delaying germination until conditions are suitable for seedling establishment and survival or by spreading germination in time (Pausas et al., 2022; Zhang et al., 2022). Despite the importance of life-history events, dormancy patterns, and the cascading effects of these adaptive traits on plant establishment and survival (Gremer et al., 2020b), our understanding of global patterns in the eco-evolutionary correlations among adaptive traits related to germination success received scant attention. Ecologists have not yet reached a conclusion on the different roles of ecologically crucial life-history traits such as seed mass and have not fully unravelled the underlying factors governing this critical functional trait. It has been recognized that germination traits are of considerable significance to plant evolutionary adaptations and distribution (Zhang et al., 2022). To coin the hypothesis on the importance of seed traits in plant adaptation, we must attempt to determine how varying environments shape seed-related traits by taking the coordinated evolution of these traits into account.

Seed mass and dormancy can temporally and spatially vary, depending on species and geographical origin, leading to heterogeneity in environments. The mechanism behind variable seed dormancy has been defined as bet-hedging, suggesting a strategy of spreading germination over time to reduce risk of failure by taking advantage of erratic favourable conditions (Philippi, 1993). The trade-off between seed size and dormancy is well-documented (Baskin and Baskin, 2014; Volis and Bohrer, 2013). Chen et al. (2020) reported that there might have been coordinated evolution between seed mass and dormancy, with the selective interaction having positive effects on geometric mean fitness. They found that small-seeded species exhibit high levels of seed dormancy, implying that seed dormancy is an evolutionarily stable strategy of plant species in response to short spells of suitable conditions (Rubio de Casas et al., 2017). Furthermore, Rees (1996) noted that heavy seeds tend to exhibit lower levels of dormancy in comparison to their smaller counterparts. These previous observations which indicate that heavy seeds have a decreased likelihood of exhibiting dormancy may be congruent with enhanced establishment success and reduced susceptibility to herbivory (Rees, 1996). However, seed mass is also imperative in explaining seed dispersal syndromes, with large seeds being maladaptive for long-distance dispersal (Thomson et al., 2011). The reduced dispersal capacity experienced by large-seeded species can be compensated to a certain extent by other traits, including seed release height and dispersal appendages (Chen et al., 2020). Given the importance of seed dormancy and its potential interaction with seed mass, to date, many aspects of the relationship between seed mass and seed dormancy remain unclear.

Seed mass is an important ecological trait that provides a reliable link between germination characteristics and plant establishment (Verdú and Traveset 2005; Adler et al. 2014). Many aspects of a plant species’ ecological strategy are impacted by seed size, including survival rate and seed dispersal (Leishman et al., 2000). Seed size plays a vital role especially in early stages of the plants life cycle, the fate of seedlings and post-germination processes (Simons and Johnston, 2000; Moles and Westoby, 2004ab; Susko & Cavers, 2008; Larios et al., 2014). Seed size, similar to other functional traits, affects population persistence and post-germination traits, and has been shown to be under strong natural selection (Larios et al., 2014; Larios and Venable., 2018). Studies on the implications of the variation in seed size have indicated that being larger can result in producing larger and more vigorous seedlings than smaller ones (Geritz et al., 2018), putting large-seeded species at a competitive advantage by preventing the growth of seedlings that emerge later. It has been shown that the fitness advantage of larger seeds over smaller ones is related to a higher likelihood of avoiding unfavorable conditions by lower persistence in the soil seed bank (Leishman et al., 2000). Furthermore, Bohrer et al. (2008) illustrated that smaller seeds may be dispersed longer distances relative to larger seeds, which may decrease the competition from the mother plant and other neighbouring plants. Notwithstanding prior evidence suggesting a positive correlation between seed mass and seed dispersal syndrome, an alternative investigation highlights that the association between plant height and seed dispersal distance is more robust than that of seed mass (Thomson et al., 2011).

Having small seeds is tightly linked to seed survival in the soil seed bank (Leishman et al., 2000) due to higher likelihood of burial and poor emergence from deep soil (Xia et al., 2016). It has been shown that small seeds are more dependent on light (Jankowska-Blaszcuk and Daws., 2007; Xia et al., 2016). This dependency on light is considered an adaptation for small-seeded species to induce germination when seeds come closer to the soil surface and are able to capture gap detection signals. In contrast, large-seeded plant species rely more often on thermal signals, such as daily temperature fluctuations, enabling them to emerge even when buried deeply (Ghersa., 1992; Pearson et al., 2002; Xia et al., 2016). Together, these studies show that the relative importance of light requirement for germination is directly related to seed mass. However, it is crucial to consider the impact of the interaction between light requirements, seed dormancy and seed mass on germination traits, as seed dormancy may change the sensitivity of seeds to their light environment (Baskin and Baskin, 2014). Although much attention has been given to the interaction between seed mass and light requirement (Milberg et al., 2000; Xia et al., 2016), a comprehensive study on the relationship among seed mass and light requirement and its interaction with seed dormancy and life form is lacking (Flores et al., 2011).

When considering germination timing as an important early life-cycle trait, characteristics related to germination, such as germination speed, must be included, since flowering plants are able to germinate at widely variable velocities (Kadereit et al., 2017). It is believed that germination speed, which is controlled by morphological and physiological features of seeds (Nonogaki et al., 2010), is an adaptive characteristic that evolves in response to variation in environmental conditions, climate and life-history strategies (Vandelook et al., 2012a; Kadereit et al., 2017). Studies have shown that species with larger embryos can germinate rapidly, suggesting that this might be an adaptive strategy to climatic or biotic conditions for germination and seedling establishment (Vandelook et al., 2012ab). Evolution has driven germination speeds to extremes in several species growing in very dry or saline habitats, resulting in germination within a matter of hours or a few days, depending on species and dormancy type (Parsons, 2012; Parsons et al. 2014). Although previous studies clarified the relationship between seed mass and life-history traits and some functional traits, it remains unclear whether the empirical studies are in alignment with theoretical evidence (Moles et al., 2007; Nordern et al., 2009), assuming that large-seeded species are able to germinate quickly and to higher percentage compared to species with small seeds.

Databases providing information on multiple ecologically meaningful traits now contain data for large number of species throughout the world. This implies that we are now able to study adaptive traits at a global scale. To study the interplay of seed mass and morphological and physiological traits related to germination behavior of plant species with a global distribution, we used advanced analytical methods to address more specifically the following hypotheses: (1) large-seeded species show increased performance as compared to small-seeded ones in terms of germination and seedling emergence, (2) some seed traits might show coordinated evolution adapted to specific environments, (3) due to the trade-off between seed mass and seed dormancy, plants may show different strategies to ensure survival, (4) germination responses to varying seed mass may differ among various dormancy types, and (5) germination traits related to seed mass are phylogenetically conserved. We expect that large-seeded species germinate faster and to a higher percentage than small-seeded ones, while high levels of seed dormancy are expected to be present in small seeds. Moreover, we expect that some post-germination traits, including seedling emergence and seedling emergence rate show coordinated evolution with seed mass since these traits are thought to be adaptive and vulnerable to environmental conditions, leading to increased selection pressures.

## 2. Material and methods

### 2.1. Study selection

Data on seed size and germination were collected from literature using the Web of Science database, with the objective of aggregating relevant studies covering the period from 1977 to 2022. Since numerous studies are available, the articles were classified according to the ‘relevance option’ as provided by the Web of Science, and all the studies that fit the criteria mentioned below were used in the meta-analysis. This process covered the following terms: seed size + germination (183), seed mass + germination (47), seed weight + germination (54), seed size + seedling emergence (36) seed mass + seedling emergence (8) and seed weight + seedling emergence (11). We targeted studies having data on four critical traits as follows: *Germination percentage, Inverse time to germination – germination rate, Seedling emergence* and *Inverse time to seedling emergence – emergence rate.* The study inclusion requirements were as follows: (1) species or populations were tested for the variation in seed size in relation to the above-mentioned traits, and studies were laid out under at least two different treatments, (2) treatments included testing germination characteristics of differing seed size of a single species, (3) studied species were endemic (or well adapted) to the study site (to make sure that species maintain an evolutionary record of the collection area), and (4) studies with consistent seed size measure (mg. or any convertible unit except for studies reporting seed length as this unit was not convertible into mg) were taken into account. In order to ensure consistency in reporting seed mass across various studies, we adopted a standardized approach. We followed the methodology outlined by Vandelook et al. (2008) and Rosbakh and Poschlod (2015) to determine seed mass for each species. These methods involved the measurement of seed mass by randomly selecting and weighing 100 seeds, subsequently calculating the mean weight per seed. In cases where studies deviated from these established methods, we transformed their data into standardized values. Based on these criteria, 339 studies were found which provided germination kinetics of nearly 1877 plant species. To obtain the suitable number of studies required for observing main effects, Rosenberg’s fail-safe-numbers method was employed, which is a method estimating the average of null results that should be added to the observed outcomes to adjust the significance level (i.e., p-value; data not shown) (Koricheva et al., 2013).

### 2.2. Data collection and plant classification

Plant species were categorized into three groups based on their growth form, specifically woody species (shrub and trees), herbs (Moles et al., 2007). This classification was adopted to expand upon a previous extensive investigation into the correlation between seed mass and germination traits, with the inclusion of additional species and case studies (Norden et al., 2009). Regarding the classification of plant species based on their dormancy type, a comprehensive database compiled by Baskin and Baskin (2014) was employed in the current study. Using this database, we categorized plant species into one of five distinct seed dormancy types: physiological, physical, morphological, morphophysiological, and non-dormancy. In order to investigate the possible interplay of dormancy level (classification) and seed mass, the preceding database was also employed to differentiate between dormant and non-dormant seeds.

### 2.3. Co-evolution of seed mass and light response

We extracted information pertaining to germination responses to light from the dataset initially established by Milberg et al. (2000) encompassing 54 distinct species. We added additional data sourced from 30 scholarly articles spanning the period from December 2000 to January 2023, which added 50 previously unaccounted species to the dataset. Furthermore, we integrated a cohort of species (previously documented by Grime et al., 1981), We used Web of Science (WOS) database to extract these articles by using the keywords “seed mass and light response” and “seed mass and light requirement”.

We incorporated RLG (relative light germination) into this study by computing relative light germination as suggested by Milberg et al. (2000):

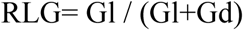

where Gl = the germination percentage in light, and Gd = the germination percentage in darkness. We considered relative values preferable to germination percentages since seed batches differed in their dormancy level. RLG represents a range of values varying from 0 (germination only in darkness) to 1 (germination only in light).

Given the recognized variability of germination-related traits and light responsiveness across different life forms (Maleki et al., 2022), in our study we included five distinct life forms: annual forbs, annual grasses, perennial forbs, perennial grasses, and shrubs, as recommended by Grime et al. (1981).

### 2.4. Ratio of embryo versus seed mass

Data on embryo length to seed length ratio and embryo to seed surface ratio were obtained mainly from pictures or drawings available in the literature dealing with the internal morphology of seeds (e.g. Martin, 1946; Watson and Dallwitz, 1992; Royal Botanic Gardens Kew, 2021) and from measurements on living material (Vandelook *et al*., 2012, 2021). Measurements were taken on transverse sections of ripe seeds, which had maximal embryo surface area. The embryo to seed length ratio (EL/SL) was calculated by measuring total embryo length and dividing this by the length of the seed, measured from the inside of the seed coat, along the longest axis. The embryo to seed surface area (ES/SS) was calculated by dividing the embryo area by the area of the embryo plus endosperm and perisperm. In the case of curved or coiled embryos, the lengths of different segments were summed. In total, we managed to obtain EL/SL data for 422 species and ES/SS data for 425 species.

### 2.5. Statistical approaches

#### 2.5.1 Meta-analysis approach

A meta-analysis of primary data, which is a method analysing original data from different studies together, was performed (Koricheva et al., 2013; Borenstein et al., 2021). We used the data within studies published to synthesize a comprehensive conclusion at the eco-evolutionary role of seed mass in controlling seed germination traits.

To examine whether seed mass has effects on germination traits a meta-analysis of primary data was performed by running binomial phylogenetic generalized mixed models with Bayesian estimation using the Markov chain Monte Carlo (MCMC) approach (Hadfield, 2010), following previous studies that used the same method for related traits (Fernández-Pascual et al., 2021; Carta et al., 2022). This approach effectively accounts for multiple observations per species/studies while considering phylogenetic relationships among different species (Carta et al., 2022; Garamszegi, 2014). This approach also enabled us to account for between-study variation owing to different scales and units used for germination traits and seed mass. In this analysis, final germination percentage, germination rate, final seedling emergence percentage and seedling emergence rate were treated as responses variable while the fixed effects for our models (the predictors) included seed mass gradient (different values of seed mass reported within each study for each germination trait); see section 2.1 and 2.2 for more detail on study selection and data collection. In our models, weak informative Gelman priors in combination with parameter expanded priors for the random effects and residual variance fixed to 1 (Hadfield, 2010). The model involved executing each procedure with four separate chains, each including 500,000 MCMC (Markov Chain Monte Carlo) steps. The initial 50,000 steps of each chain were disregarded as a ’burn-in’ phase, and the data was taken every 100 steps (de Villemereuil & Nakagawa, 2014). The outputs derived from these four chains consisting of 500,000 MCMC were then combined to obtain parameter estimates. The mean of these parameter output and their 95% credible intervals (CIs) were computed from the merged posterior distributions. To see if the parameters are significant, we examined these CIs; parameters were considered insignificant if their CIs included zero. The explanatory aspect of our analysis was evaluated through calculating the conditional R^2^ (i.e. the amount of variation accounted for by both the fixed (traits) and random factors (the phylogeny)) and the marginal (i.e. the amount of variation accounted for by the fixed factors only) using the approach described by Nakagawa et al. (2013)

#### 2.5.2. Multivariate ordination method

To test whether was a relationship between seed mass and germination traits, principal component analysis (PCA) implemented within the R environment (FACTOMINER package) was employed (Le et al., 2008; Rosbakh et al., 2023). We first quantified the potential relationship for each growth form through fitting univariate models. Then, predicted probabilities of the relationship for each trait to which they belong to were used in the multivariate ordination.

#### 2.5.3. Binary approach

To answer the long-standing question of how seed mass interacts with differing levels of seed dormancy, we used Bayesian logistic phylogenetically informed generalized mixed models, following Rosbakh et al. (2023). In the models, we treated seed dormancy (the response variable) as binary data. Here, 0 and 1 refer to the absence or presence of seed dormancy(0= seed mass of non-dormant seeds; 1= seed mass of dormant seeds), respectively. This binary classification refers to the primary distinction in dormancy states relevant to our research question. For the fixed effects in our models, seed mass was treated as the predictor. To directly compare the response/predictor effects, we centred and scaled the variance of all the variables to be unit. To assess this relationship, the model was fitted in two distinct manners: firstly, through comparing the seed mass of non-dormant versus dormant seeds (specifically water-permeable versus impermeable for physical dormancy), to see how seed mass might affect the likelihood of dormancy; and secondly, by examining this correlation within each specific dormancy class separately. We aimed to understand potential variations in the seed mass- dormancy relationship across different dormancy types, suggesting that different classes might respond differently to variations in seed mass. As a result, each observation in the analysis was a species with specific seed dormancy type and corresponding seed mass value.

#### 2.5.4 Phylogenetic regression

To analyse the relation between RLG and seed mass and between embryo to seed size ratio and germination rate we applied a phylogenetic regression with ML procedure, whereby the phylogenetic signal and regression model were estimated simultaneously. As a measure of the phylogenetic signal, we applied Pagel’s *λ* (Pagel, 1999). When *λ* equals zero, related taxa are not more similar than expected by chance and the trait is evolving as in star-like phylogeny (Pagel, 1999). In such a scenario, phylogenetic correction becomes redundant. Significant phylogenetic signal or clumping of trait states on the phylogenetic tree occurs when *λ* > 0, meaning taxa are more similar than expected by chance. When *λ* = 1, the trait is evolving following a constant variance random walk or BM model. If 1 > *λ* > 0, traits are less similar among species than expected from their phylogenetic relationships, but more similar than expected by chance. Analyses were performed using the APE (Paradis, 2006) and nlme (Pinheiro *et al*., 2009) packages in R.

## 3. Results

### 3.1. Relative proportion of seed mass

The final database consisted of 1877 plant species representing 140 families for which we had data on germination traits and seed mass covering all the tree growth forms included (Fig S1). The species included in this study had a seed mass range varying from 0.0001 mg to 10000 mg. Seeds of herbs mostly occurred within a range varying from 0.01 mg to 100 mg, with more than 80% of species concentrated in the 0.1 mg to 100 mg range (Fig 1ac). Most tree species tend to have a seed mass between 0.1 mg and 10000 mg, with the average value of 1.01 mg (Fig1a). The number of tree species is normally distributed across seed mass gradient, with more than 15% of species showing heavy seeds. Shrubs are mostly concentrated in the 0.01 mg to 10 mg range of seed masses, which is very similar to the trend reflected in herbs.

**Figure 1.**
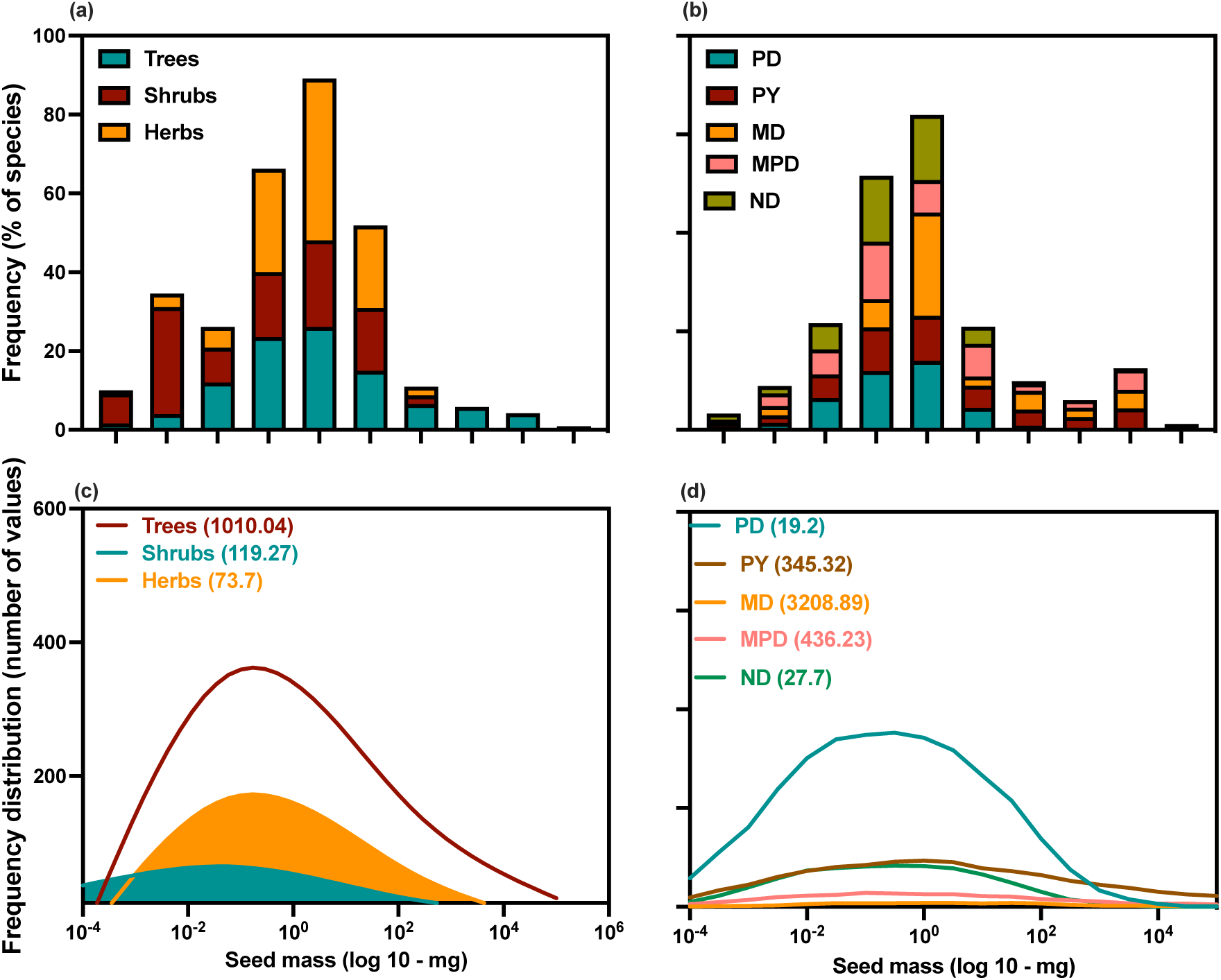
The frequency distribution of seed mass. Panels coded by (a) the frequency percentage of species grouped into different life forms, including trees, shrubs, and herbs across seed mass gradient (on a log scale) (b) the frequency percentage of species grouped into different dormancy types across seed mass gradient. (c) indicates the frequency distribution of seed mass for three different plant growth forms: trees, shrubs, and herbs, (d) shows the frequency distribution of seed mass for five different dormancy classes. The height of each curve represents the number of values collected and incorporated into the database, not the seed mass of each category. Instead, the magnitude of seed mass (mg) of each category is shown on the X-axis. Values in the parenthesis indicate the average value of seed mass in mg.

Seed mas ranges varied also among seed dormancy type, with some groups like PD and ND having a more pronounced frequency in the lower seed mass categories (Fig 1bd). MD, MPD and PY showed heavier seeds relative to PD and ND, with more than 50% of these species having a more pronounced frequency in the higher seed mass categories (Fig 1bd).

### 3.2. Univariate analyses

The univariate models predicting the distribution of seed mass and germination traits revealed that both pre-and post-germination traits may show contrasting responses when taking growth form into account. Except for trees which showed a negative relationship, suggesting that small tree seeds germinate to the highest percentage (Fig 2; Table 1). Herbs and shrubs demonstrated a positive correlation between seed mass and germination percentage. The general trend of seed germination percentage (pooled data) was also positively related to seed mass.

**Figure 2.**
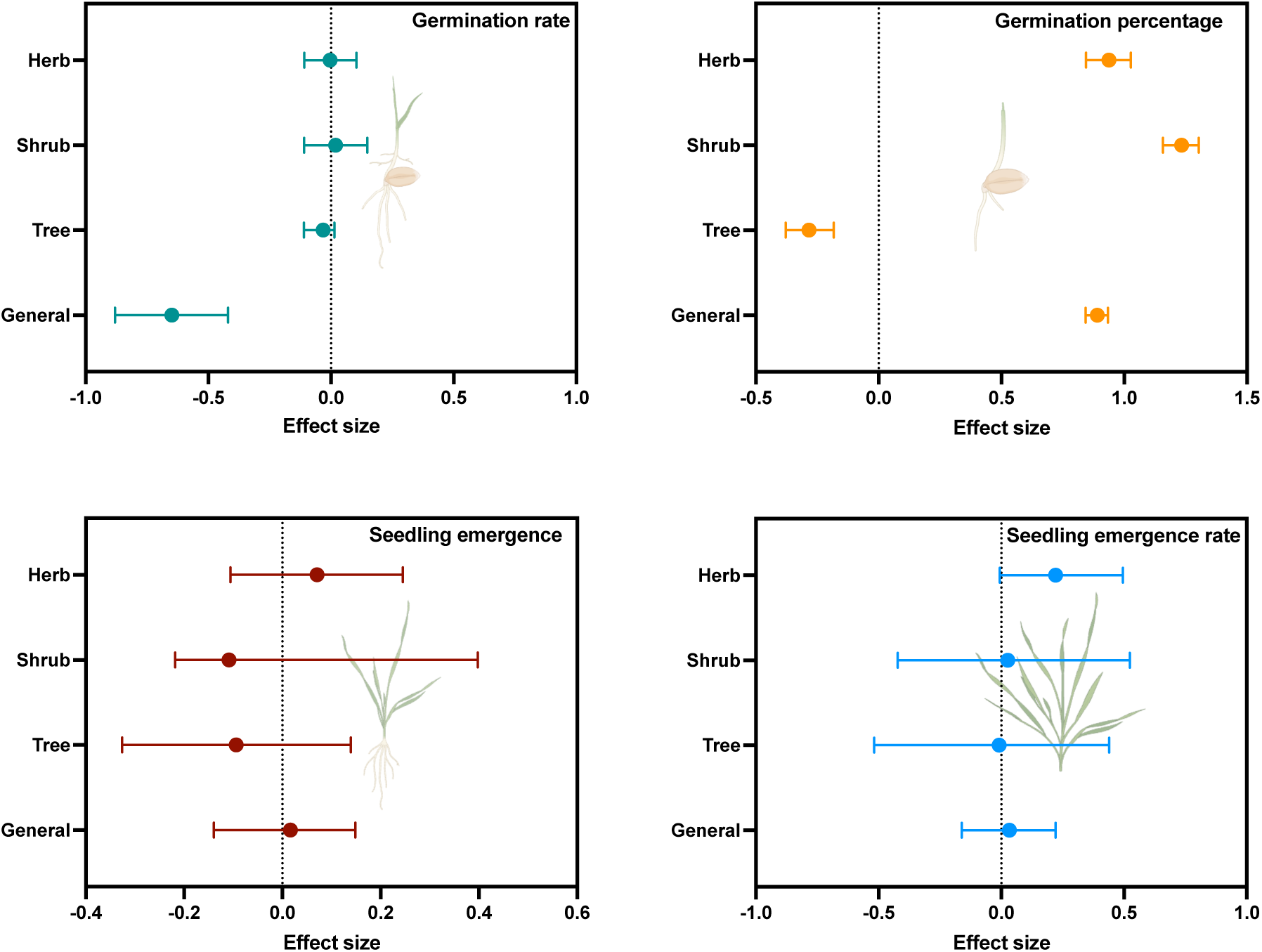
Seed mass effects on germination traits of three distinct growth forms according to the binomial phylogenetic mixed models with Bayesian estimation (MCMCglmms). Closed symbols represent the posterior means of the interaction effect size. Horizontal bars indicate the 95% credible intervals. Dashed vertical line denotes zero effect. On the top panel, germination rate and germination refer to speed of germination and percentage of germination respectively. On the bottom panel, seedling emergence and seedling emergence rate indicate seedling emergence percentage and speed, respectively.

**Table 1.**
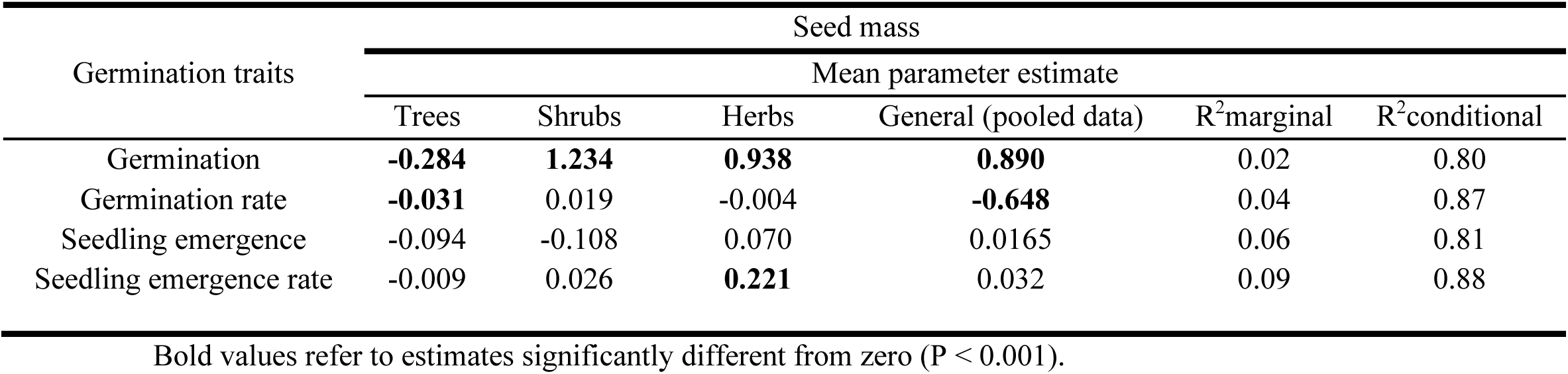
Effects of seed mass on germination traits across different growth forms using a global database as estimated by Bayesian logistic phylogenetically informed generalized mixed models.

When pooling data together (general trend), germination rate was negatively correlated with seed mass. Within growth forms, this relationship varied, with trees being negatively correlated with seed mass (**-0.031**), while shrubs (0.019) and herbs (-0.004) showed a non-significant but positive relationship between germination rate and seed mass (Fig 2; Table 1).

No significant relation between seed mass and seedling emergence was detected, nor for the pooled data, nor for the individual life-forms (Fig 2; Table 1). while germination rate of herbs was positively correlated with seed mass. The seedling emergence rate was positively related to seed mass for the pooled data (0.032) and for herbs (**0.221**) in particular, although this relationship was only significant for herbs (Fig 2; Table 1).

Our models exhibited comparatively weak the explanatory power, as the marginal R^2^ was rather limited in all the models. The cumulative amount of variance accounted for by our models was highest in seedling emergence rate (R^2^ = 0.09) followed by seedling emergence (0.06) and germination rate (0.04). The cumulative amount of variance accounted for by our models was lowest in germination percentage (0.02). Interestingly, the conditional R^2^, the major contributor to phylogenetic relatedness, was higher in all models.

### 3.3. The correlation between seed dormancy type and seed mass

The relationship between seed dormancy and seed mass was highly dependent on dormancy type, with MD, MPD and PY being significantly positively related to seed mass, suggesting that seeds of species with MD, MPD and PY are heavier (Fig 3 Table 2). A negative relation with seed mass was observed for species with PD (Fig 3 Table 3)

**Figure 3.**
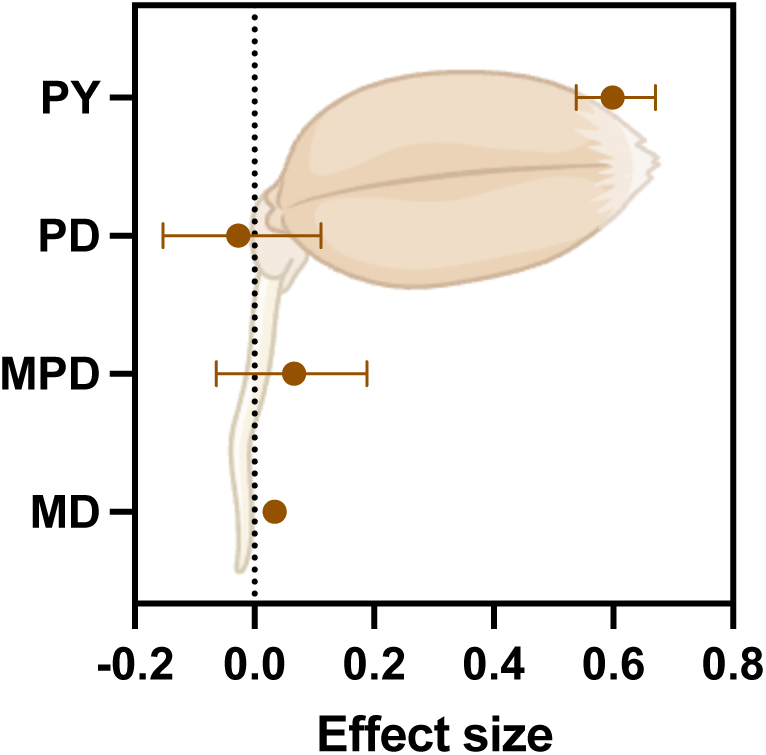
Seed mass effects on dormancy levels for four dormancy types according to the binary approach. Closed circles show the posterior means of the interaction effect size. Horizontal bars represent the 95% credible intervals. Dashed vertical line denotes zero effect. MD, morphological; MPD morphophysiological dormancy; PD, physiological dormancy; PY, physical.

**Table 2.**
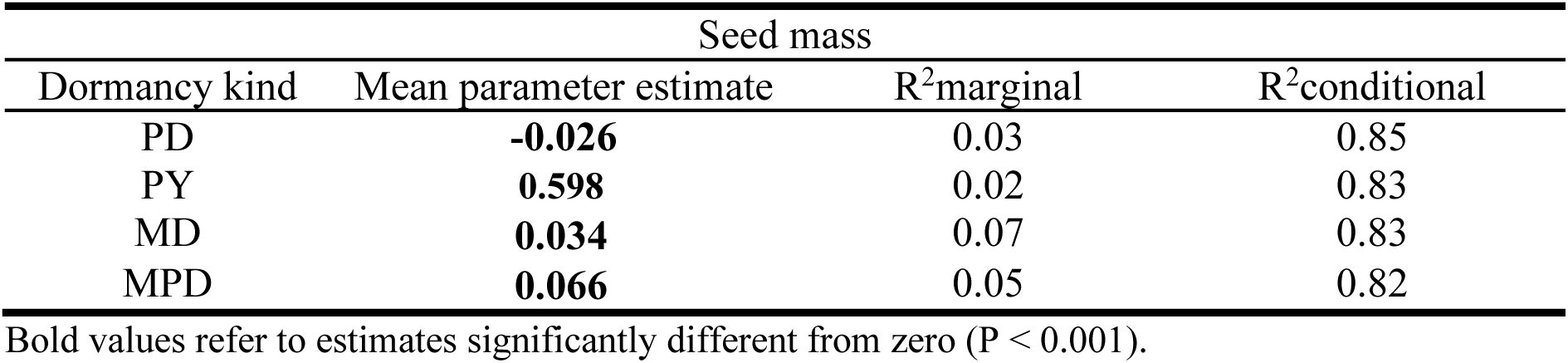
Effects of seed mass on dormancy levels across different growth forms as estimated by Bayesian logistic phylogenetically informed generalized mixed models.

| Dormancy kind | Seed mass |  |  |
| --- | --- | --- | --- |
|  | Mean parameter estimate | R<sup>2</sup>marginal | R<sup>2</sup>conditional |
| PD | **-0.026** | 0.03 | 0.85 |
| PY | **0.598** | 0.02 | 0.83 |
| MD | **0.034** | 0.07 | 0.83 |
| MPD | **0.066** | 0.05 | 0.82 |

Bold values refer to estimates significantly different from zero ( $P < 0.001$ ).

### 3.4. Correlation between seed mass and light response (RLG)

Light response was significantly negatively correlated with seed mass (Fig 4a). Small-sized seeds thus showed higher germination in light, while the more seed mass increased, light germination decreased.

**Figure 4.**
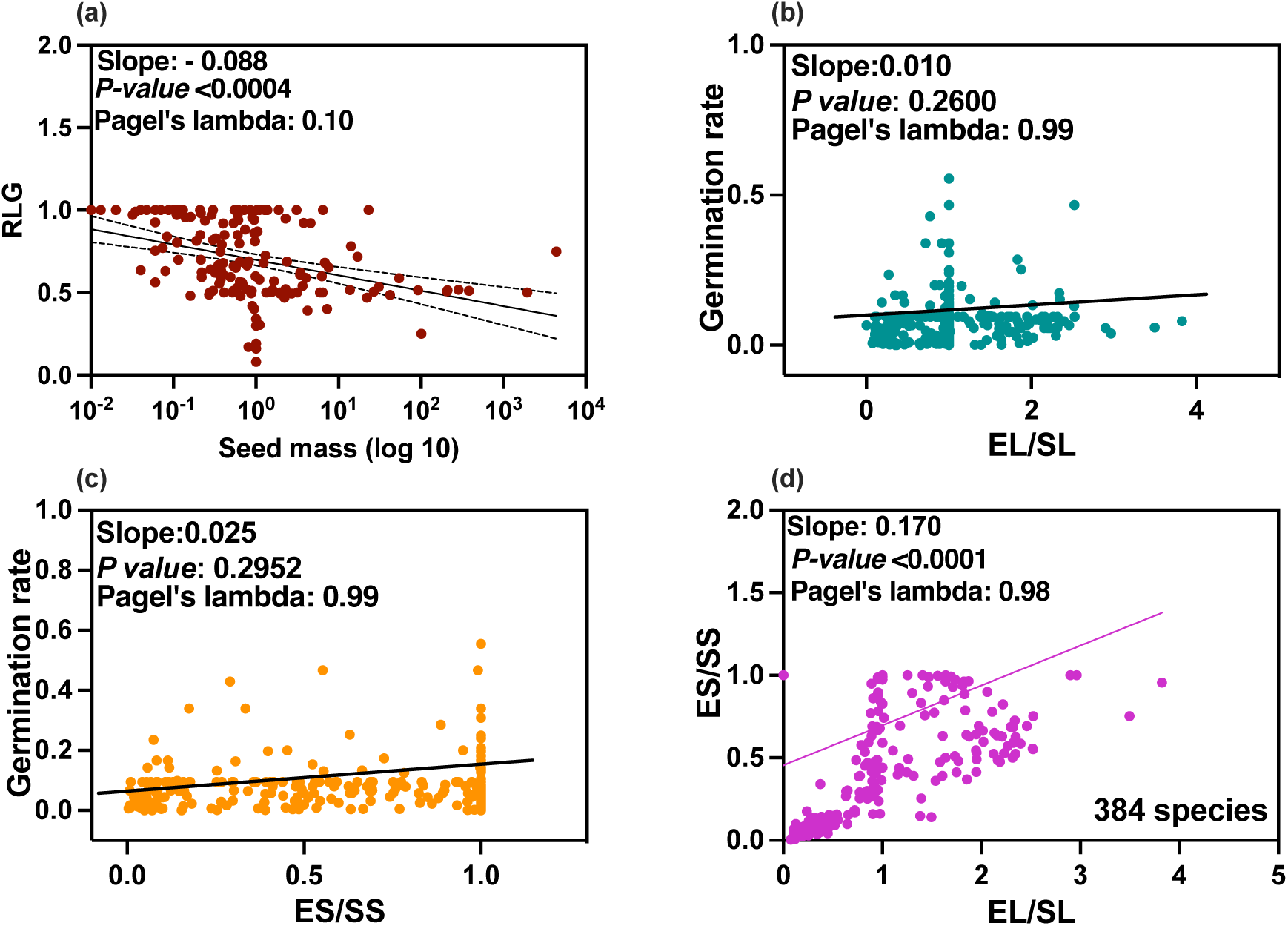
(a) Correspondence between seed mass and relative light germination. Phylogenetic regression of of relative light germination (RLG) on seed mas). (b-d) Phylogenetic egression of germination rate on EL/SL (embryo length/seed length) and ES/SS (embryo size/seed size). Fitted lines were estimated via phylogenetic regression approach (Revell., 2010).

Notably, a parallel pattern emerged within the spectrum of plant life forms (Figure S2), reflecting the previously observed trend across all species encompassed in the analysis (Figure 4a). Of the five diverse plant life forms integrated into our study, annual and perennial forbs exhibited the most pronounced negative slopes (P-value: < 0.0001 and 0.0021, respectively), showing values of -0.13 and -0.17 respectively (Pagel’s λ: 0.11 and 0.12, respectively). This indicates that the light responsiveness of forbs displays a relatively diminished reliance on light intensity compared to the responses of other life forms, including shrubs and annual grasses. Moreover, the phylogenetic signal λ observed for RLG in different growth forms was close to 0.

Pagel’s λ was close to 0 when considering the relationship between RLG and seed mass, suggesting that correlation between seed mass and RLG is not phylogenetically conserved at the family level (Fig 4).

### 3.5. Relationship between germination rate and EL/SL and ES/SS

A positive but non-significant (P < 0.096) correlation was found between germination rate and EL/SL, with a phylogenetic regression slope of 0.042 (Fig 4b). A similar trend was also reflected in the relationship between germination rate and ES/SS, with the phylogenetic regression slope of 0.046, showing a smooth but non-significant correlation between germination rate and ES/SS (Fig 4c; P = 0.244). Interesting, phylogenetic regression showed that both EL/SL, ES/SS and their correlation with seed mass is phylogenetically conserved (Pagel’s λ: 0.99; Fig 4).

### 3.6. Multivariate ordination

Two principal components summarized the variation in seed mass, germination traits and growth form by taking 58% of the total variation into account (Fig 5). 33% of variance was explained by the first component (PC1). PC1 was mainly loaded on seedling emergence, seed mass and seedling emergence rate (Fig 5). Thus, the first principal separates species with heavier seed mass and promoted seedling emergence rate and percentage grouping into shrub and herb category. 24.9% of the total variation observed was explained by the second component (PC2). Germination rate and germination percentage were the main contributing variables on axis 2 which are classified mostly into trees and herbs (Fig 5).

**Figure 5.**
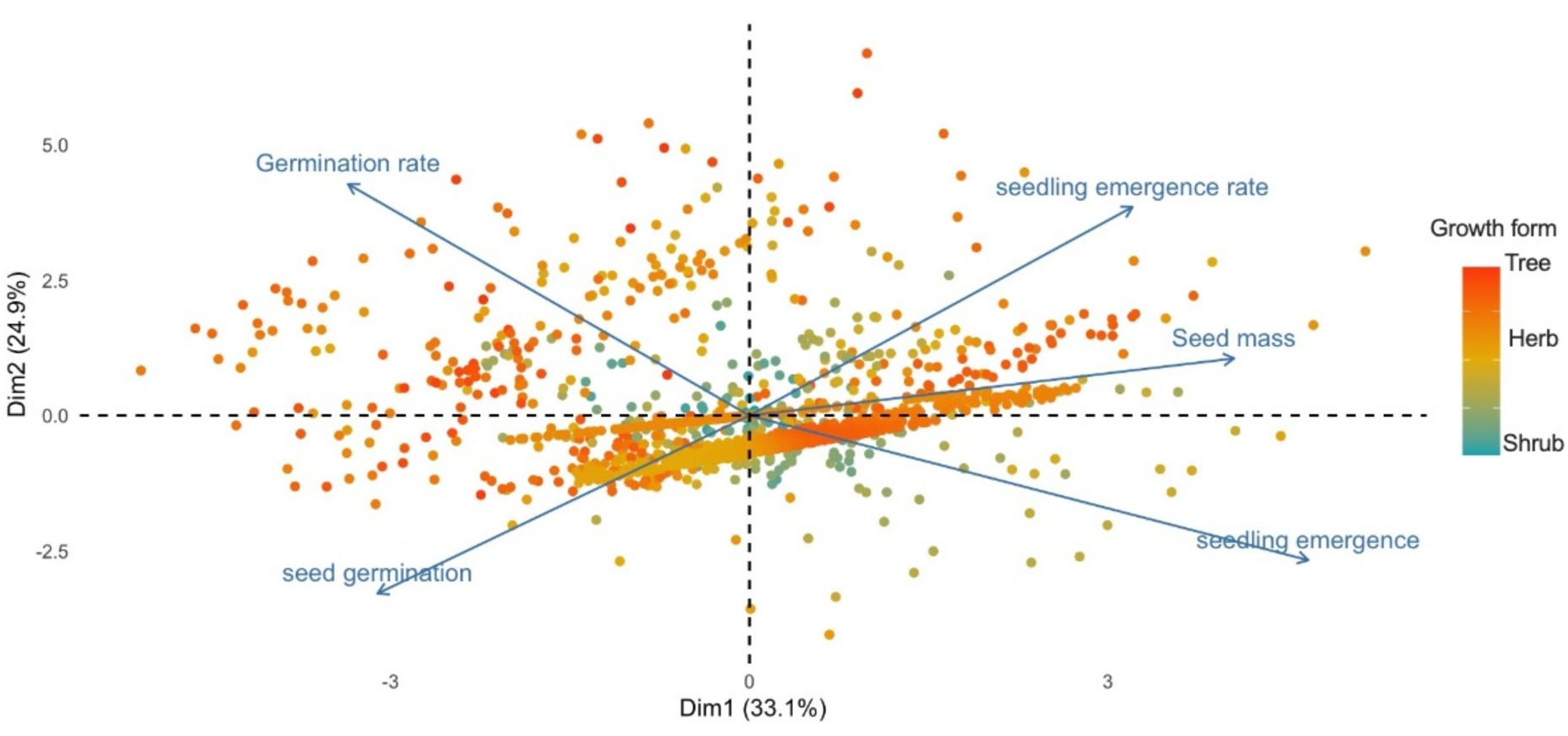
Principal component analysis (PCA) ordination representing the relationships among germination traits and seed mass. Each closed point refers to a species coloured by growth form.

## 4. Discussion

The relationship between germination traits and seed mass varied considerably among growth forms. Trees revealed a consistent negative relationship between seed mass and germination rate and seedling emergence rate. Our findings highlight that responses of germination traits to seed mass gradients are complex and highly dependent on growth forms and probably also environmental conditions experienced by plants. Seed mass variation also depended on seed dormancy type. Seeds with PD was negatively correlated with seed mass while seeds with PY, MD and MPD showed a positive relationship with seed mass.

### 4.1. Correlates of seed mass across growth forms and dormancy type

Our results show that the magnitude of the increase in variation of seed mass was related to both dormancy type and growth forms, which is consistent with previous studies exploring the seed mass gradient in different growth forms (Moles & Westoby, 2003l Moles et al 2007). Classifying seed mass into life form classes and plotting frequency histograms of seed mass distributions revealed that the magnitude of variation in seed mass varies among species and life forms (in agreement with Carta et al., 2022). It has been well documented that the latitudinal gradient explain much of the global variation (> 51%) in seed mass (Moles et al., 2007), but this is the first report of the magnitude of seed mass variation within different dormancy types. Previous studies have also reasoned that global variation in seed mass might be an important driver of species distribution with more than 46% of variation in species distribution accounted for seed mass, seed mass variability and seed dispersal mode (Moles et al., 2007; Chen et al., 2022). Variation in seed mass can explain geographical ranges within which plants can thrive, leading to the different ecological strategies that have evolved in plants (Chen et al., 2022). Moreover, empirical studies have reported that there is a trade-off between seed mass and seed dormancy (e.g. Chen et al., 2020). This relation between seed mass and dormancy can result from the trade-offs between seed mass and dormancy incurred by morphological constraints and the allocation cost of making larger seeds (Smith and Fretwell, 1974). Moreover, the preceding studies implied that selection might have operated on these two traits, either by increased dispersal or increased dormancy, meaning that a relationship between dormancy and seed mass cannot be excluded (Philippi, 1998; Volis and Bohrer., 2012). The fact that seed mass has been under strong natural selection is well-known (Larios et al., 2014; Larios and Venable., 2018), which in turn may prove that some environmental conditions may favor specific strategies which can be either heavier seeds or increased dormancy level.

### 4.2. Correlation between seed mass and germination

Seed germination is a crucial step in the life cycle of plants. Much attention has been given to studies examining various species that are different in germination strategies in response to environmental filters (Gremer et al., 2020a,b; Maleki et al., 2024). Our results suggest that germination and post-germination traits are affected by the seed mass gradient, but also that this relationship is complex and highly dependent on growth form. According to the theoretical perspective, it was believed that seed mass interacted with germination so that species with heavier seeds seem superior in their final germination proportion (Venable and Brown, 1988). However, the germination fraction might be contingent on seed viability and seed dormancy (Baskin and Baskin., 2014; Carta et al., 2022). Our findings revealed that this might not be empirically true because species with different growth forms and evolutionary history may adopt various strategies and more importantly, coordination of traits and biological trade-offs may alter the ways plants respond. To coin the preceding discussion, we found that germination is positively correlated with seed mass in herbs and shrubs, but a negative relationship was observed for trees. The negative relationship between seed mass and germination suggests that trees with small seeds may germinate to the highest percentage, which may be the result of a specific strategy maintained by trees as perennial plants. For example, this may be explained by the potential competition from neighbouring and maternal plants that exists in the forest understory which in turn may select for long-distance dispersal or increased level of seed dormancy, requiring smaller and lighter seeds to easily disperse or to enter deeper soil layers to form a soil seed bank. Additionally, most tree species exhibit PD (Rosbakh et al., 2023) which favors delayed germination and requires dormancy breaking agents (i.e, stratification and after ripening) to overcome dormancy, leading to cueing of germination to periods suitable conditions (Escobar et al., 2021; Maleki et al., 2022; Just et al., 2023). Interestingly, this hypothesis is consistent with our findings about small seeds of species with PD showing higher levels of seed dormancy (Fig 3; Table 2). As discussed later, several external factors can also control the interaction between seed mass and germination, including a light requirement for germination and alternating temperature cycles (Bond et al., 1999; Pons, 2000).

### 4.2. Eco-evolutionary relationship between seed mass and dormancy

Unlike theoretical evidence suggesting that seed mass is independent of seed dormancy (Volis and Bohrer, 2013), we found that a relationship exists between seed mass and seed dormancy, although this relationship is complex and varies among dormancy classes. Our results showed that the relationship between seed mass and dormancy highly depends on dormancy types, with PD being negatively correlated with seed mass while PY, MD and MPD are positively correlated with seed mass. This positive response might result from the embryo size or embryo length to seed length ratio and embryo to seed surface ratio. In seeds with MD and MPD, a small or undifferentiated embryo impedes germination until the embryo grows to a certain extent. Species with MD and MPD are typically grow in shaded environments, which also explains their higher seed mass (Baskin and Baskin, 2014; Vandelook et al. 2012).

A higher seed size we observed in seeds with physical dormancy is in agreement with the crypsis hypothesis invoked by Paulsen et al. (2014) to explain evolution of the water impermeable seed coat in as a predation avoidance mechanism. In previous studies it was already observed that physical dormancy is absent from lineages with predominantly small seeds (Leishman & Westoby, <u>1998</u>; Šerá & Šerý, <u>2004</u>). In contrast, in a study on seeds of 14 species with physical dormancy, it was shown that the threshold temperature for seed dormancy breaking is higher in small seeds than in larger seeds (Liyanage and Ooi, 2018). From a plant strategy perspective, this might be a mechanism for smaller seeds that respond to fire as a germination cue to avoid germination too deep in the soil, as the temperatures required to overcome physical dormancy may not be reached when seeds are buried too deep in the soil (Liyanage and Ooi, 2018). In a meta-analysis of the effects of frugivory (endozoochory) on seed germination, results indicated that germination percentage of all seed mass categories, in seeds with physical dormancy, was increased by gut passage, and the magnitude of the increase in germination percentage was higher in large seeds than in medium and small seeds (Soltani et al., 2018). This suggests that the water-impermeable seed coat provides better protection in adverse environments, enabling seeds to reach larger sizes. On the other hand, once the seed coat becomes permeable, the protective function is lost and the large seeds profit from germinating rapidly (Dalling et al. 2011).

In the current study, the relationship between seed mass and seed viability was not considered due to dearth of information on this interaction, which calls for future studies on the understanding of the regulatory roles of seed viability in driving seed mass. Furthermore, a few studies reported that no close relationship exists between seed dormancy and seed viability (Thompson et al., 2003; Long et al., 2015).

### 4.3. Correlation between seed mass and germination rate

Seeds are able to germinate at various speeds. The time required to shift from the beginning of imbibition to radicle protrusion may vary depending on the seed coat thickness and environmental factors underlying germination (Baskin and Baskin, 2014). Contradictory results have been reported regarding the relationship between seed size and germination rate; in some of them germination rate is significantly and negatively correlated with seed mass (e.g. Norden et al., 2009) while in other reports a temperature dependent positive and significant relationship was found between seed size and germination rate (e.g. Vandelook et al., 2012a). The advantage of meta-analysis in the present study is that we can provide a global and general result based on all the available information, that as a result, small seeds had a higher germination rate and this relationship was statistically significant (Table 1; general trend) Although the general trend (pooled data) was negative positive, showing smaller seeds germinate faster, but this relationship seems to be complex as it varies among growth forms with shrubs showing a positive correlation, meaning that larger seeds are faster than small ones.

The negative correlation between seed mass and rate of germination is in agreement with empirical observations (Norden et al., 2009). In contrast theoretical studies emphasized that larger seeds may germinate faster and this argument is contingent on the survivorship of large-sized seeds due to a higher post-dispersal predation, indicating that a high pressure imposed by post-dispersal predation might promote rapid germination in larger seeds (Janzen 1971; Louda 1989). The preceding discussion may also explain the positive correlation between seed mass and germination rate observed for shrubs. Furthermore, a study elaborating on data for hundreds of plant species with worldwide distribution suggested that there is no evidence of a negative relationship between seed mass and seed survivorship (Moles et al., 2003). While a previous study reported no positive correlation between seed mass and germination speed (Norden et al., 2009), we found that germination rate indeed accelerates with increasing seed mass in shrubs which highlights the complexity of the correlation between seed mass and germination rate. This observation aligns with theoretical predictions and supports the notion that seed mass and germination rate have coevolved as an alternative mechanism to cope with unpredictable environments (Venable and Brown, 1988; Kadereit et al., 2017). One strength of our study is the global scope, as we conducted a comprehensive analysis encompassing various habitats, climates, and species with diverse life-forms, dormancy types, and phylogenetic relationships which suggest that the relationship between germination rate and seed mass is probably species-specific, suggesting a complex eco-evolutionary history.

Our results often align with the findings of other studies which highlighted the negative relationship between seed mass and germination rate. For example, Murali (1997) reported that germination rate of small seeds from non- tropical aeras where light availability is not limited was faster. However, it has been also shown that germination rate of tropical trees is negatively correlated with seed mass, which might originate from developmental and physiological constraints (Norden et al. 2009). One of the factors underlying the preceding observed patterns might be physical barriers, such as a thicker seed coat, which is of great importance because larger seeds may want to invest more in protection against predation (Tiansawat et al., 2014).

### 4.5. Correlation between seed mass and seedling emergence and rate

Much attention has been given to the correlation between seed mass and seedling emergence (Singh and Saxena., 2009). Studies have shown that seedling emergence is associated with germination rate and timing (Mishra et al., 2014; Cao et al., 2018). Seedling emergence rate may not be sensitive to seed dormancy because it is not just a function of the germination process but is also influenced by other factors, such as root growth, shoot growth, or resource availability (Gardarin et al., 2016). Furthermore, the rate of seedling emergence could be more related to the underlying developmental processes, such as the timing of organ initiation or tissue differentiation, rather than seed dormancy status (Vandelook and Van Assche 2008).

The observed relationship between seed mass and germination rate has been contradictory in different studies, with some studies showing small-seeded species germinate rapidly and other studies discussing the general model that large-seeded species should germinate more rapidly (Venable and Brown, 1988; Rees, 1994). Our results showed that, except for trees, seedling emergence rate becomes faster as the seed mass increases and this relationship was statistically positive when fitting model on pooled data. The reason behind the fact that seedling emergence rate of larger seeds is faster than small ones is that large-seeded species are prone to higher post-dispersal seed predation and herbivory, meaning that faster emergence of larger seeds favors survival through establishing prior to being destroyed or grazed by herbivores (Dalling et al., 2011; Cao et al., 2018). More interestingly, studies have reported that the reason given for higher seedling emergence of larger seeds is that seedlings grown from large seeds have higher amounts of inherent carbon metabolic rates as well as seed resource consumption rates, giving rise to more vigorous seedlings (Green and Juniper 2004, Quero et al. 2007). However, other studies arrived at a different conclusion (Norden et al., 2009). For example, no relationship between seed mass and seedling survival was reported (Leishman., 1999). These contradictory conclusions may result from a lack of studies on the relationship between seed mass and seedling emergence. Also in our study, less data were available on seedling emergence and seedling emergence rate within scientific literature, which calls for further research on these ecologically meaningful traits.

Furthermore, studies have revealed that the level of dormancy varies across species (Baskin and Baskin, 2014) and even among different populations of the same species (Maleki et al., 2022), resulting in divergent requirements for dormancy loss in seeds. This ecological rationale provides insights into the underlying factors contributing to the variation in seed mass observed both within and among species. In this regard, studies have indicated that superiority of larger seeds compared with smaller ones might be related to embryo viability and higher carbohydrate and protein content (González-Rodríguez et al. 2011; Bu et al., 2019). Our results showed a positive correlation between embryo size and germination rate. A positive correlation between germination traits and increasing seed mass has been reported in herbs and trees (Seltmann et al. 2007; Dunlap & Barnett 1983). It has been shown that larger seeds are likely to be more successful than smaller ones in establishment due to vigorous seedlings produced by larger seeds (Baraloto et al., 2005; Bruun & Ten Brink, 2008; Volis and Bohrer., 2012; Domic et al., 2020). Many factors may exert effects on the relationship between seed mass and germination traits, including sowing depth, litter and herbaceous cover (Weller, 1985; Wulff, 1986; Gulmon, 1992; Rebollo et al., 2001; Dalling & Hubbell, 2002). By virtue of higher germination percentages and higher rate of seedling emergence, larger seeds are better than small- and medium-sized seeds. The preceding reports are in agreement with the findings of our study showing that larger seeds enable plants to gain a competitive advantage but this relationship might not be true in all situations and depends on other eco-evolutionary factors. This ecological reasoning offers new insights into how plants leverage a broad spectrum of traits to optimize their germination strategies, by either increasing seed mass or seed dormancy. In the face of unpredictably changing environments, rapid germination becomes crucial for plants to exploit favorable conditions, thereby implying reduced bet-hedging. Conversely, bet-hedging serves as a survival strategy by spreading germination over an extended period, increasing the likelihood of seed survival to the following year compared to simultaneous germination of all seeds (Gremer and Venable., 2014; Ten Brink et al., 2020). Additionally, although larger seeds and the resulting vigorous seedlings can better tolerate unsuitable environments but the risk of germinating immediately even if conditions are not suitable become lower as compared to smaller seeds.

### 4.6. Correlated evolution of the germination response to light and seed mass

Light requirements for germination have been studied for decades (Milberg et al., 2000; Baskin and Baskin, 2014; Xia et al., 2016). Smaller seeds more often require light to complete germination, while large-seeded species seem less dependent on light (Grime et al., 1981; Milberg et al., 2000). The reason given for this is that large-seeded species can emerge successfully from deeper layers of soil than light can penetrate (del Arco et al., 1995). In addition, light serves as a gap detection mechanism, which benefits small-seeded pioneer species that establish after a disturbance in the vegetation (Vandelook et al. 2008). Our results indicate that as seed mass increases, seeds become less dependent on light to complete germination. These findings are consistent with the hypothesis proposed by Milberg et al. (2000) and the empirical framework of Grime et al. (1981), indicating that light as an environmental signal triggering germination becomes less important in species with relatively large seeds. However, it is worth noting that RLG differs considerably and that a high amount of the variation in RLG is explained by seed mass. Furthermore, many species germinated better in light than darkness (points below regression line in Fig 4). Since the slope is negative, it is assumed that light consistently seems to be less important in large-seeded species. In agreement with our findings, studies have shown that there is a trade-off between seed mass and light responses (Milberg et al., 2007; Rubio de Casas et al., 2017; Santana et al., 2020).

### 4.7. The correlation between EL/SL and ES/SS and germination rate

A positive relation between relative embryo size (expressed as EL/SL or ES/SS) with germination has been observed in both Apiaceae (Vandelook et al. 2012) and Amaranthaceae (Vandelook et al. 2021). This relation has been explained by the fact that the transfer of nutrients from endosperm or perisperm to the growing seedling is time consuming. In our global study, we have shown that the global relation between germination speed and relative embryo size is non-significant. This agrees with the results of Verdu (2006) who equally showed that there was no relationship between relative embryo size and germination speed using a global data-set. The reason for this discrepancy may be that different plant families exhibit widely different dormancy mechanisms and seed morphologies, that may mask potential relationships between relative embryo size and germination speed.

## Conclusion

The dispersal of seeds from the maternal plants exposes them to a range of biotic and abiotic threats to their survival, such as predation and environmental hazards. Given the significance of early life-history strategies in ensuring plant survival and population persistence, germination is considered a critical stage in the plant life cycle. Seed mass, being a crucial life-history trait, is expected to have a profound impact on germination success, plant survival, and reproduction. Our findings provide evidence of a positive correlation between increasing seed mass and pre- and post-germination traits, further highlighting the importance of this life-history trait in the overall success of plants. Seed dormancy, as an important trait regulating adaptation, seems to have coevolved with seed mass gradient as an alternative to confront unpredictable environments. Interestingly, the sensitivity of germination to light decreases with increasing seed mass, suggesting that the light requirement for germination is primarily associated with lighter seeds. This implies that germination responses to light might have evolved independently of the general light requirement, possibly representing a distinct strategy for regulating germination adaptation. A substantial portion of variability in germination characteristics remains unaccounted for, indicating the presence of additional factors that contribute to the broad interspecific variation in the correlation between germination traits and seed mass observed in plants. In order to enhance our comprehension of the evolutionary and ecological factors that influence germination traits, it is imperative to conduct comprehensive studies that incorporate both community-level and multivariate approaches.

## Supporting information

Supplementary file-phylogeny

## Author contributions

Keyvan Maleki assembled and managed the database, analyzed the data and drafted the manuscript, conceived the original idea. Filip Vandelook, Arne Saatkamp and Elias Soltani commented on the data analyses and revised the manuscript. Koroush Maleki, Siavash Heshmati and Filip Vandelook collected and extracted the data from the literature. All authors approved the final version.

## References

Adler, P. B., Salguero-Gómez, R., Compagnoni, A., Hsu, J. S., Ray-Mukherjee, J., Mbeau-Ache, C., & Franco, M. 2014. Functional traits explain variation in plant life history strategies. Proceedings of the National Academy of Sciences, 111: 740–745. doi: 10.1073/pnas.1315179111.

Bohrer, G., Katul, G. G., Nathan, R., Walko, R. L., & Avissar, R. 2008. Effects of canopy heterogeneity, seed abscission and inertia on wind-driven dispersal kernels of tree seeds. Journal of Ecology, 96: 569–580. 10.1111/j.1365-2745.2008.01368.x.

Bond WJ, Honig M, Maze KE. 1999. Seed size and seedling emergence: an allometric relationship and some ecological implications. Oecologia 120: 132– 136. 10.1007/s004420050841.

Bu, H. Y., Zhang, Y. M., Zhao, D., Wang, S. Y., Jia, P., Qi, W., … & Wang, X. J. 2019. The evolutionary correlation associated with seed mass and altitude on nutrient allocation of seeds. Seed Science Research, 29: 38–43. 10.1017/S0960258518000387.

Buckley, L. B, Kingsolver, J. G. 2012. Functional and phylogenetic approaches to forecasting species’ responses to climate change. *Annual Review of Ecology*, Evolution and Systematics, 43: 2012. 10.1146/annurev-ecolsys-110411-160516.

Baskin, C. C, Baskin, J. M. 2014. Seeds: Ecology, biogeography, and, evolution of dormancy and germination. Academic Press/Elseiver.

Baraloto, C., Forget, P. M., Goldberg, D. E. 2005. Seed mass, seedling size and neotropical tree seedling establishment. Journal of Ecology, 93: 1156–1166. 10.1111/j.1365-2745.2005.01041.x.

Bruun, H. H., & Ten Brink, D. J. 2008. Recruitment advantage of large seeds is greater in shaded habitats. Ecoscience, 15: 498–507. 10.2980/15-4-3147.

Cao, S., Liu, K., Du, G., Baskin, J. M., Baskin, C. C., Bu, H., Qi, W. 2018. Seedling emergence of 144 subalpine meadow plants: Effects of phylogeny, life cycle type and seed mass. Seed Science Research, 28: 93–99. 10.1017/S0960258518000028.

Carta, A., Mattana, E., Dickie, J., Vandelook, F. 2022. Correlated evolution of seed mass and genome size varies among life forms in flowering plants. Seed Science Research, 32: 46–52. 10.1017/S0960258522000071.

Chen, S. C., Poschlod, P., Antonelli, A., Liu, U., Dickie, J. B. 2020. Trade-off between seed dispersal in space and time. Ecology Letters, 23: 1635–1642. 10.1111/ele.13595.

Chen, K., Burgess, K. S., He, F., Yang, X. Y., Gao, L. M., Li, D. Z. 2022. Seed traits and phylogeny explain plants’ geographic distribution. Biogeosciences, 19: 4801–4810. 10.5194/bg-19-4801-2022.

Dalling, J. W., Hubbell, S. P. 2002. Seed size, growth rate and gap microsite conditions as determinants of recruitment success for pioneer species. Journal of Ecology, 557-568. 10.1046/j.1365-2745.2002.00695.x.

De Frenne, P., Graae, B. J., Rodríguez-Sánchez, F., Kolb, A., Chabrerie, O., Decocq, G., … Verheyen, K. 2013. Latitudinal gradients as natural laboratories to infer species’ responses to temperature. Journal of Ecology, 101: 784–795. 10.1111/1365-2745.12074.

Del Arco, M. S., Torner, C., Quintanilla, C. F. 1995. Seed dynamics in populations of *Avena sterilis ssp. ludoviciana*. Weed Research, 35: 477–487. DOI:10.1111/J.1365-3180.1995.TB01645.X.

Dunlap JR and Barnett JP 1983. Influence of seed size on germination and early development of loblolly pine (Pinus taeda L.) germinants. Journal of Forest Research 13: 40–44. 10.1139/x83-006.

de Villemereuil, P., Nakagawa, S. 2014. General quantitative genetic methods for comparative biology. Modern phylogenetic comparative methods and their application in evolutionary biology: concepts and practice, 287-303. DOI: 10.1111/j.1420-9101.2009.01915.x.

Domic, A. I., Capriles, J. M., Camilo, G. R. 2020. Evaluating the fitness effects of seed size and maternal tree size on *Polylepis tomentella* (Rosaceae) seed germination and seedling performance. Journal of Tropical Ecology, 36: 115–122. 10.1017/S0266467420000061.

Dalling JW, Hubbell SP. 2002. Seed size, growth rate and gap micrositeconditions as determinants of recruitment success for pioneer species. Journal of Ecology 90: 557–568. 10.1046/j.1365-2745.2002.00695.x.

Dalling, J. W., Davis, A. S., Schutte, B. J., Elizabeth Arnold, A. 2011. Seed survival in soil: interacting effects of predation, dormancy and the soil microbial community. Journal of Ecology, 99: 89–95. 10.1111/j.1365-2745.2010.01739.x.

Escobar, D. F. E., Rubio de Casas, R., Morellato, L. P. C. 2021. Many roads to success: different combinations of life-history traits provide accurate germination timing in seasonally dry environments. Oikos, 130: 1865–1879. 10.1111/oik.08522.

Fernández-Pascual, E., Carta, A., Mondoni, A., Cavieres, L. A., Rosbakh, S., Venn, S., … Jiménez-Alfaro, B. 2021. The seed germination spectrum of alpine plants: a global meta-analysis. New phytologist, 229: 3573–3586. 10.1111/nph.17086.

Flores, J., Jurado, E., Chapa-Vargas, L., Ceroni-Stuva, A., Dávila-Aranda, P., Galindez, G., … & Pritchard, H. W. (2011). Seeds photoblastism and its relationship with some plant traits in 136 cacti taxa. Environmental and Experimental Botany, 71(1), 79–88. 10.1016/j.envexpbot.2010.10.025.

Gardarin, A., Coste, F., Wagner, M. H., Dürr, C. 2016. How do seed and seedling traits influence germination and emergence parameters in crop species? A comparative analysis. Seed Science Research, 26: 317–331. 10.1017/S0960258516000210.

Geritz, S., Gyllenberg, M., Toivonen, J. 2018. Adaptive correlations between seed size and germination time. Journal of mathematical biology, 77: 1943–1968. 10.1007/s00285-018-1232-z.

González-Rodríguez, V., Villar, R., & Navarro-Cerrillo, R. M. 2011. Maternal influences on seed mass effect and initial seedling growth in four Quercus species. Acta Oecologica, 37: 1–9. 10.1016/j.actao.2010.10.006.

Green, P. T., Juniper, P. A. 2004. Seed–seedling allometry in tropical rain forest trees: seed mass-related patterns of resource allocation and the ‘reserve effect’. Journal of Ecology, 92: 397–408. https://oi.org/10.1111/j.0022-0477.2004.00889.x.

Gremer, J. R., Venable, D. L. 2014. Bet hedging in desert winter annual plants: optimal germination strategies in a variable environment. Ecology letters, 17: 380–387. 10.1111/ele.12241.

Gremer, J. R., Chiono, A., Suglia, E., Bontrager, M., Okafor, L., Schmitt, J. (2020a). Variation in the seasonal germination niche across an elevational gradient: the role of germination cueing in current and future climates. American journal of botany, 107: 350–363. 10.1002/ajb2.1425.

Gremer, J. R., Wilcox, C. J., Chiono, A., Suglia, E., Schmitt, J. 2020b. Germination timing and chilling exposure create contingency in life history and influence fitness in the native wildflower Streptanthus tortuosus. Journal of Ecology, 108: 239–255. 10.1111/1365-2745.13241.

Grime, J. P. 1977. Evidence for the existence of three primary strategies in plants and its relevance to ecological and evolutionary theory. The american naturalist, 111: 1169–1194.

Grime, J. P., Mason, G., Curtis, A. V., Rodman, J., Band, S. R. 1981. A comparative study of germination characteristics in a local flora. The Journal of Ecology, 1017-1059. DOI:10.2307/2259651.

Ghersa, C. M., Arnold, R. B., Martinez-Ghersa, M. A. 1992. The role of fluctuating temperatures in germination and establishment of Sorghum halepense. Regulation of germination at increasing depths. Functional Ecology, 460-468. 10.2307/2389403.

Garamszegi, L. Z. (Ed.). 2014. Modern phylogenetic comparative methods and their application in evolutionary biology: concepts and practice. Springer. Doi/ 10.1007/978-3-662-43550-2.

Gulmon, S. L. 1992. Patterns of seed germination in Californian serpentine grassland species. Oecologia, 89: 27–31. 10.1007/BF00319011.

Hashemirad, S., Soltani, E., Darbandi, A.I. Alahdadi, I., 2023. Cold stratification requirement to break morphophysiological dormancy of fennel (*Foeniculum vulgare Mill*.) seeds varies with seed length. Journal of Applied Research on Medicinal and Aromatic Plants, 35:100465. 10.1016/j.jarmap.2023.100465.

Tung Ho, L. S., Ané, C. 2014. A linear-time algorithm for Gaussian and non-Gaussian trait evolution models. Systematic biology, 63: 397–408. 10.1093/sysbio/syu005.

Hadfield, J. D. 2010. MCMC methods for multi-response generalized linear mixed models: the MCMCglmm R package. Journal of statistical software, 33: 1–22. 10.18637/jss.v033.i02.

Jin, Y., Qian, H. 2019. V. PhyloMaker: an R package that can generate very large phylogenies for vascular plants. Ecography, 42: 1353–1359. DOI: 10.1111/ecog.04434.

Just, M., Cross, A. T., Lewandrowski, W., Turner, S. R., Merritt, D. J., Dixon, K. 2023. Seed dormancy alleviation by warm stratification progressively widens the germination window in Mediterranean climate Rutaceae. Australian Journal of Botany, 71:55–66. doi:10.1071/BT22076.

Koricheva, J., Gurevitch, J., Mengersen, K. (Eds.). 2013. Handbook of meta-analysis in ecology and evolution. Princeton University Press.

Kadereit, G., Newton, R. J., Vandelook, F. 2017. Evolutionary ecology of fast seed germination—a case study in Amaranthaceae/Chenopodiaceae. Perspectives in Plant Ecology, Evolution and Systematics, 29: 1–11. 10.1016/j.ppees.2017.09.007.

Larios, E., & Venable, D. L. 2018. Selection for seed size: The unexpected effects of water availability and density. Functional Ecology, 32: 2216–2224. 10.1111/1365-2435.13138.

Larios, E., Búrquez, A., Becerra, J. X., Lawrence Venable, D. 2014. Natural selection on seed size through the life cycle of a desert annual plant. Ecology, 95: 3213–3220. DOI:10.1890/13-1965.1.

Leishman, M. R. 1999. How well do plant traits correlate with establishment ability? Evidence from a study of 16 calcareous grassland species. The New Phytologist, 141:487-496. 10.1046/j.1469-8137.1999.00354.x.

Leishman, M. R., Wright, I. J., Moles, A. T., Westoby, M. (2000). The evolutionary ecology of seed size. In Seeds: the ecology of regeneration in plant communities (pp. 31–57). Wallingford UK: CABI publishing. DOI: 10.1079/9780851994321.0031.

Leishman, M.R., Westoby, M. Jurado, E. 1995 Correlates of seed size variation — a comparison among five temperate floras. Journal of Ecology, 83: 517–529. 10.2307/2261604.

Leishman, M. R., Westoby, M. 1998. Seed size and shape are not related to persistence in soil in Australia in the same way as in Britain. Functional Ecology, 12: 480–485. 10.1046/j.1365-2435.1998.00215.x.

Liyanage, G.S. Ooi, M.K., 2018. Seed size-mediated dormancy thresholds: a case for the selective pressure of fire on physically dormant species. Biological Journal of the Linnean Society, 123:135–143. 10.1093/biolinnean/blx117.

Lê, S., Josse, J., Husson, F. 2008. FactoMineR: an R package for multivariate analysis. Journal of statistical software, 25: 1–18. 10.18637/jss.v025.i01.

Maleki, K., Baskin, C. C., Baskin, J. M., Kiani, M., Alahdadi, I., Soltani, E. 2022. Seed germination thermal niche differs among nine populations of an annual plant: A modeling approach. Ecology and evolution, 12: e9240. 10.1002/ece3.9240.

Maleki, K., Soltani, E., Seal, C. E., Colville, L., Pritchard, H. W., Lamichhane, J. R. 2024. The seed germination spectrum of 486 plant species: A global meta-regression and phylogenetic pattern in relation to temperature and water potential. Agricultural and Forest Meteorology, 346: 109865. 10.1016/j.agrformet.2023.109865.

Milberg, P., Andersson, L., & Thompson, K. 2000. Large-seeded spices are less dependent on light for germination than small-seeded ones. Seed science research, 10: 99–104. 10.1017/S0960258500000118.

Mishra, Y., Rawat, R., Rana, P. K., Sonkar, M. K., Mohammad, N. 2014. Effect of seed mass on emergence and seedling development in Pterocarpus marsupium Roxb. Journal of forestry research, 25: 415–418. 10.1007/s11676-014-0469-7.

Moles, A. T., Westoby, M. 2004b. What do seedlings die from and what are the implications for evolution of seed size?. Oikos, 106: 193–199. 10.1111/j.0030-1299.2004.13101.x.

Moles, A.T., Ackerly, D.D., Tweddle, J.C., Dickie, J.B., Smith, R., Leishman, M.R., Mayfield, M.M., Pitman, A., Wood, J.T. Westoby, M., 2007. Global patterns in seed size. Global ecology and biogeography, 16:109–116. DOI: 10.1111/j.1466-8238.2006.00259.x.

Moles, A.T. & Westoby, M. 2003. Latitude, seed predation and seed mass. Journal of Biogeography, 30: 105–128. 10.1046/j.1365-2699.2003.00781.x.

Moles, A.T., Ackerly, D.D., Webb, C.O., Tweddle, J.C., Dickie, J.B. Westoby, M. 2005a. A brief history of seed size. Science, 307: 576–580. DOI: 10.1126/science.1104863.

Moles, A.T., Ackerly, D.D., Webb, C.O., Tweddle, J.C., Dickie, J.B., Pitman, A.J. & Westoby, M. 2005. Factors that shape seed mass evolution. Proceedings of the National Academy of Science USA, 102: 10540–10544. doi: 10.1073/pnas.0501473102.

Moles, A. T., Westoby, M. 2004a. Seedling survival and seed size: a synthesis of the literature. Journal of Ecology, 92:372–383. 10.1111/j.0022-0477.2004.00884.x.

Martin, A. C. 1946. The comparative internal morphology of seeds. The American Midland Naturalist, 36: 513–660. 10.2307/2421457.

Murali, K. S. 1997. Patterns of seed size, germination and seed viability of tropical tree species in southern India 1. Biotropica, 29: 271–279. 10.1111/j.1744-7429.1997.tb00428.x.

Nakagawa, S., Schielzeth, H. 2013. A general and simple method for obtaining R2 from generalized linear mixed-effects models. Methods in ecology and evolution, 4: 133–142. 10.1111/j.2041-210x.2012.00261.x.

Nonogaki, H., Bassel, G. W., Bewley, J. D. 2010. Germination—still a mystery. Plant Science, 179: 574–581. 10.1016/j.plantsci.2010.02.010.

Norden, N., Daws, M. I., Antoine, C., Gonzalez, M. A., Garwood, N. C., Chave, J. 2009. The relationship between seed mass and mean time to germination for 1037 tree species across five tropical forests. Functional Ecology, 203-210. 10.1111/j.1365-2435.2008.01477.x.

Parsons, R. F. 2012. Incidence and ecology of very fast germination. Seed Science Research, 22: 161–167. 10.1017/S0960258512000037.

Parsons, R. F., Vandelook, F., Janssens, S. B. 2014. Very fast germination: additional records and relationship to embryo size and phylogeny. Seed Science Research, 24: 159–163. 10.1017/S096025851400004X.

Pearson, T. R. H., Burslem, D. F. R. P., Mullins, C. E., Dalling, J. W. 2002. Germination ecology of neotropical pioneers: interacting effects of environmental conditions and seed size. Ecology, 83: 2798–2807. 10.1890/0012-9658(2002)083[2798:GEONPI]2.0.CO;2.

Pons TL. 2000. Seed responses to light. In: Fenner M, ed. Seeds: the ecology of regeneration in plant communities. Wallingford, UK: CABI, 237–260. 10.1079/9780851994321.0237.

Pausas, J. G., Lamont, B. B., Keeley, J. E., Bond, W. J. 2022. Bet-hedging and best-bet strategies shape seed dormancy. The New Phytologist, 236: 1232. 10.1111/nph.18436.

Paulsen, T. R., Colville, L., Kranner, I., Daws, M. I., Högstedt, G., Vandvik, V., Thompson, K. 2013. Physical dormancy in seeds: a game of hide and seek?. New phytologist, 198: 496–503. 10.1111/nph.12191.

Philippi, T. 1993. Bet-hedging germination of desert annuals: beyond the first year. The American Naturalist, 142: 474–487.

Quero, J. L., Villar, R., Marañón, T., Zamora, R., Poorter, L. 2007. Seed-mass effects in four Mediterranean Quercus species (Fagaceae) growing in contrasting light environments. American Journal of Botany, 94:1795–1803. 10.3732/ajb.94.11.1795.

Rodrigues-Junior, A.G., Mello, A.C.M., Baskin, C.C., Baskin, J.M., Oliveira, D.M. Garcia, Q.S. 2018. Why large seeds with physical dormancy become nondormant earlier than small ones. PLoS One, 13: p.e0202038. 10.1371/journal.pone.0202038.

Rosbakh, S., Poschlod, P. 2015. Initial temperature of seed germination as related to species occurrence along a temperature gradient. Functional Ecology, 29: 5–14. 10.1111/1365-2435.12304.

Rosbakh, S., Carta, A., Fernández-Pascual, E., Phartyal, S.S., Dayrell, R.L., Mattana, E., Saatkamp, A., Vandelook, F., Baskin, J. and Baskin, C., 2023. Global seed dormancy patterns are driven by macroclimate but not fire regime. New Phytologist, 240:555–564. 10.1111/nph.19173.

Rubio de Casas, R., Willis, C. G., Pearse, W. D., Baskin, C. C., Baskin, J. M., Cavender-Bares, J. 2017. Global biogeography of seed dormancy is determined by seasonality and seed size: a case study in the legumes. New Phytologist, 214: 1527–1536. 10.1111/nph.14498.

Rees, M. 1996. Evolutionary ecology of seed dormancy and seed size. Philosophical Transactions of the Royal Society of London. Series B: Biological Sciences, 351:1299–1308. 10.1098/rstb.1996.0113.

Rebollo, S., Pérez-Camacho, L., García-de Juan, M. T., Rey Benayas, J. M., Gómez-Sal, A. 2001. Recruitment in a Mediterranean annual plant community: seed bank, emergence, litter, and intra-and inter-specific interactions. Oikos, 95: 485–495. 10.1034/j.1600-0706.2001.950314.x.

Revell, L. J. 2010. Phylogenetic signal and linear regression on species data. Methods in Ecology and Evolution, 1: 319–329. 10.1111/j.2041-210X.2010.00044.x.

Santana, V. M., Alday, J. G., Adamo, I., Alloza, J. A., Baeza, M. J. 2020. Climate, and not fire, drives the phylogenetic clustering of species with hard-coated seeds in Mediterranean Basin communities. Perspectives in Plant Ecology, Evolution and Systematics, 45: 125545. 10.1016/j.ppees.2020.125545.

Seltmann P, Leyer I, Renison D Hensen I 2007. Variation of seed mass and its effects on germination in Polylepis australis: implications for seed collection. New Forest 33:171–181. 10.1007/s11056-006-9021-8.

Simons, A. M., Johnston, M. O. 2000. Variation in seed traits of *Lobelia inflata* (Campanulaceae): sources and fitness consequences. American Journal of Botany, 87: 124–132. 10.2307/2656690.

Singh N, Saxena AK. 2009. Seed size variation and its effect on germination and seedling growth of *Jatropha curcas L*. Indian Forester, 135: 1135−1142. DOI: 10.36808/if/2009/v135i8/447.

Smith, C. C., Fretwell, S. D. 1974. The optimal balance between size and number of offspring. The American Naturalist, 108: 499–506. 10.1086/282929.

Šerá, B., Šerý, M. 2004. Number and weight of seeds and reproductive strategies of herbaceous plants. Folia Geobotanica, 39: 27–40. 10.1007/BF02803262.

Soltani, E., Baskin, C.C., Baskin, J.M., Heshmati, S. Mirfazeli, M.S. 2018. A meta-analysis of the effects of frugivory (endozoochory) on seed germination: role of seed size and kind of dormancy. Plant Ecology, 219: 1283–1294. 10.1007/s11258-018-0878-3.

Susko, D. J., Cavers, P. B. 2008. Seed size effects and competitive ability in *Thlaspi arvense L*.(Brassicaceae). Botany, 86: 259–267. 10.1139/B07-137.

Ten Brink, H., Gremer, J. R., Kokko, H. 2020. Optimal germination timing in unpredictable environments: the importance of dormancy for both among-and within-season variation. Ecology letters, 23: 620–630. 10.1111/ele.13461.

Thomson, F. J., Moles, A. T., Auld, T. D., Kingsford, R. T. 2011. Seed dispersal distance is more strongly correlated with plant height than with seed mass. Journal of ecology, 99: 1299–1307. 10.1111/j.1365-2745.2011.01867.x.

Tiansawat, P., Davis, A. S., Berhow, M. A., Zalamea, P. C., Dalling, J. W. 2014. Investment in seed physical defence is associated with species’ light requirement for regeneration and seed persistence: Evidence from Macaranga species in Borneo. PLoS One, 9: e99691. 10.1371/journal.pone.0099691.

Vandelook, F., Van Assche, J. A. 2008. Temperature requirements for seed germination and seedling development determine timing of seedling emergence of three monocotyledonous temperate forest spring geophytes. Annals of Botany, 102: 865–875. 10.1093/aob/mcn165.

Vandelook, F., Janssens, S. B., Probert, R. J. 2012. Relative embryo length as an adaptation to habitat and life cycle in Apiaceae. New Phytologist, 195: 479–487. 10.1111/j.1469-8137.2012.04172.x.

Vandelook, F., Van de Moer, D., Pajolec. L 2021. Embryo to seed length ratio and embryo to seed surface ratio data [Data set]. Zenodo. 10.5281/zenodo.5647046.

Vandelook, F., Van de Moer, D., Van Assche, J. A. 2008. Environmental signals for seed germination reflect habitat adaptations in four temperate Caryophyllaceae. Functional Ecology, 22: 470–478. 10.1111/j.1365-2435.2008.01385.x.

Vandelook, F., Verdú, M., Honnay, O. 2012. The role of seed traits in determining the phylogenetic structure of temperate plant communities. Annals of Botany, 110: 629–636. 10.1093/aob/mcs121.

Verdú, M., Traveset, A. (2005). Early emergence enhances plant fitness: a phylogenetically controlled meta-analysis. Ecology, 86: 1385–1394. 10.1890/04-1647.

Volis, S., Bohrer, G. 2013. Joint evolution of seed traits along an aridity gradient: seed size and dormancy are not two substitutable evolutionary traits in temporally heterogeneous environment. New Phytologist, 197: 655–667. 10.1111/nph.12024.

Westoby, M., Jurado, E., Leishman, M. 1992. Comparative evolutionary ecology of seed size. Trends in Ecology and Evolution, 7: 368–372. DOI: 10.1016/0169-5347(92)90006-W.

Watson, L. 1992. The families of flowering plants: descriptions, illustrations, identification and information retrieval. http://biodiversity.uno.edu/delta.htm.

Weller, S. G. 1985. Establishment of *Lithospermum caroliniense* on sand dunes: the role of nutlet mass. Ecology, 66: 1893–1901. 10.2307/2937385.

Wulff, R. D. 1986. Seed size variation in *Desmodium paniculatum*: II. Effects on seedling growth and physiological performance. The Journal of Ecology, 99-114. 10.2307/2260351.

Xia, Q., Ando, M., Seiwa, K. 2016. Interaction of seed size with light quality and temperature regimes as germination cues in 10 temperate pioneer tree species. Functional Ecology, 30: 866–874. 10.1111/1365-2435.12584.

