## Supplementary file-phylogeny for "Global patterns in the evolutionary relations between seed mass and germination traits"

### Supplementary information

#### *Phylogenetic tree:*

To generate a phylogeny of the studied species, we used the largest dated mega-tree for vascular plants (Open Tree of Life) as a backbone. An ultrametric phylogenetic tree containing the species in our data-set was then constructed using the package *V.PhyloMaker* (Jin and Qian, 2019) implemented within the R environment.

Jin, Y., & Qian, H. (2019). V. PhyloMaker: an R package that can generate very large phylogenies for vascular plants. Ecography, 42(8), 1353-1359.

### Supplementary Figures

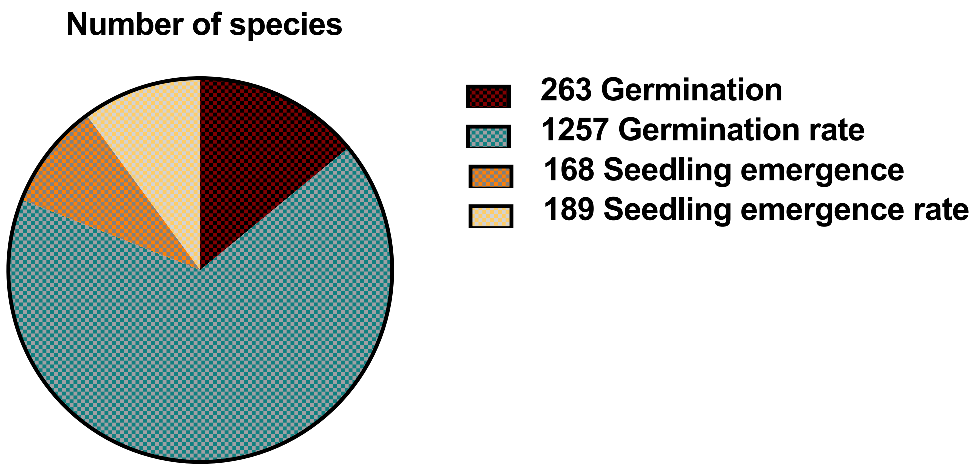


**Fig S1.** The graph presented depicts the number of species included in the analysis of traits across various plant categories and dormancy types. The figures provided in the legend correspond to the total number of species represented in the graph.

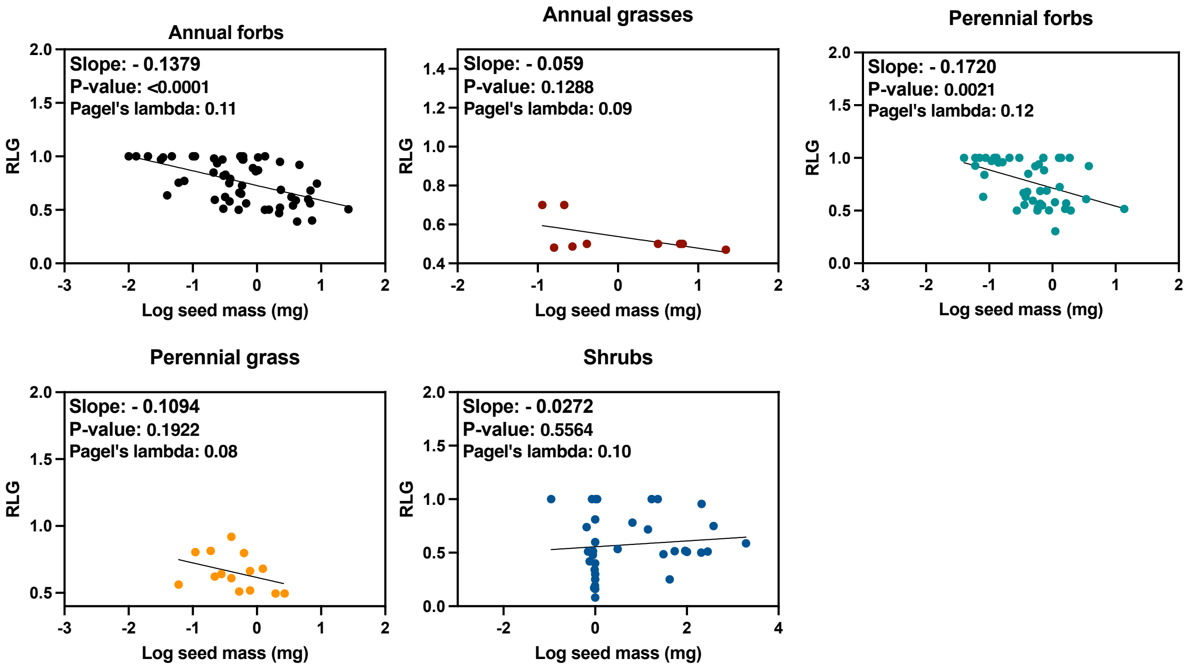


**Fig S2.** Correspondence between seed mass and relative light germination in relation to life forms. Regression of relative light germination (RLG) on seed mass. Fitted lines were estimated via phylogenetic regression approach as suggested by Revell., 2010.

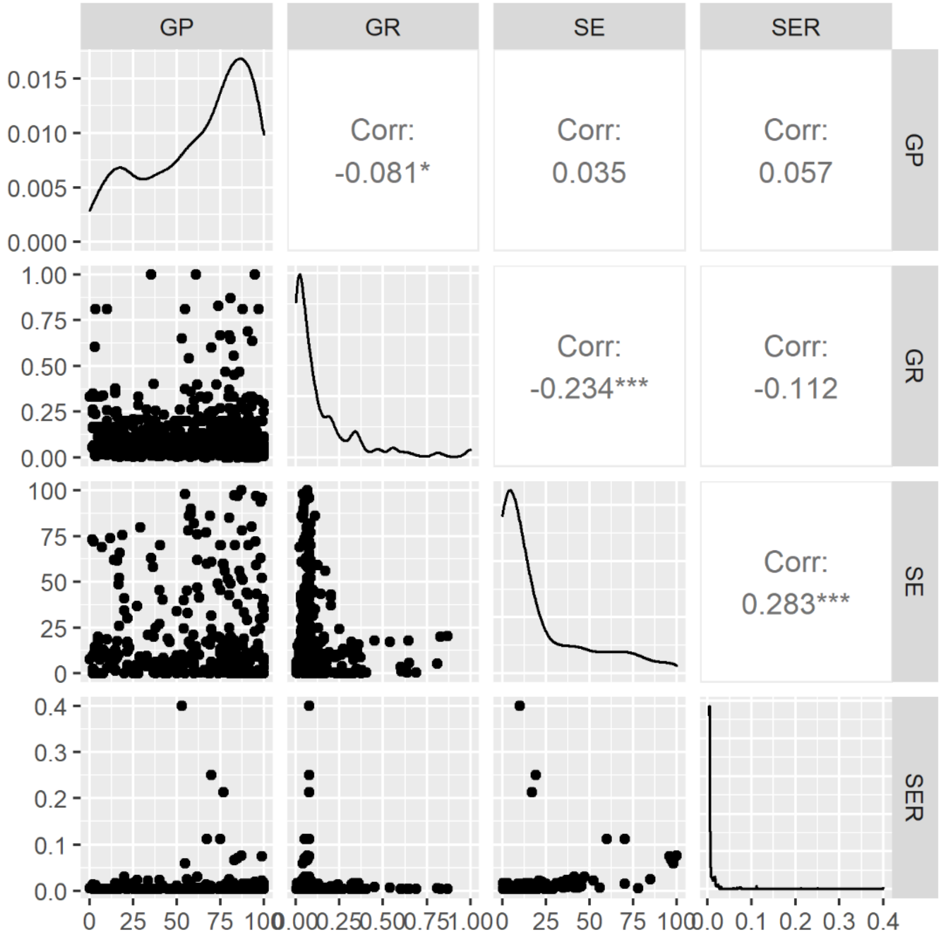


**Fig S3.** Correlation between germination traits as affected by increasing seed mass.

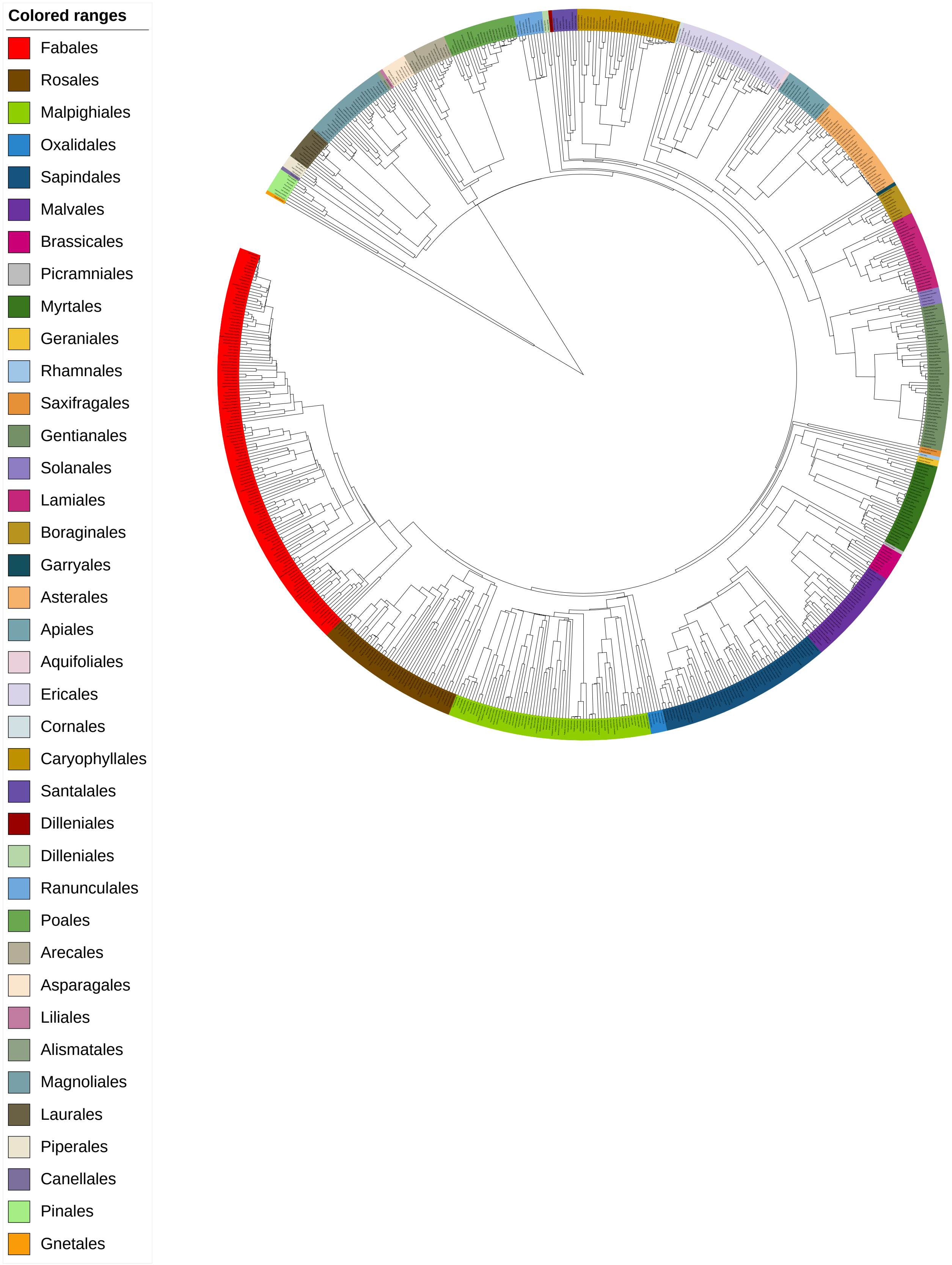
Colored ranges
Fabales
Rosales
Malpighiales
Oxalidales
Sapindales
Malvales
Brassicales
Picramniales
Myrtales
Geraniales
Rhamnales
Saxifragales
Gentianales
Solanales
Lamiales
Boraginales
Garryales
Asterales
Apiales
Aquifoliales
Ericales
Cornales
Caryophyllales
Santalales
Dilleniales
Dilleniales
Ranunculales
Poales
Arecales
Asparagales
Liliales
Alismatales
Magnoliales
Laurales
Piperales
Canellales
Pinales
Gnetales

**Fig S4.** The phylogenetic relationships of germination traits and seed masss. Coloured clades show taxonomic orders of 1877 species.
